# COInr and mkCOInr: Building and customizing a non-redundant barcoding reference database from BOLD and NCBI using a lightweight pipeline

**DOI:** 10.1101/2022.05.18.492423

**Authors:** Emese Meglécz

**Author notes:** Corresponding author: Emese Meglécz, Aix Marseille Univ, Avignon Univ, CNRS, IRD, IMBE, Chemin de la batterie des Lions, 13007 Marseille FRANCE.

## Abstract

The taxonomic assignment of metabarcoding data strongly depends on the taxonomic coverage of the reference database. Therefore, it is fundamental to access and pool data from the two major sources of COI sequences, the BOLD and the NCBI nucleotide databases, and enrich them with custom COI data, when available.

The COInr database is a freely available, easy-to-access database of COI reference sequences extracted from the BOLD and NCBI nucleotide databases. It is a comprehensive database: not limited to a taxon, a gene region, or a taxonomic resolution; therefore, it is a good starting point for creating custom databases. Sequences are dereplicated between databases and within taxa. Each taxon has a unique taxonomic Identifier (taxID), fundamental to avoid ambiguous associations of homonyms and synonyms in the source database. TaxIDs form a coherent hierarchical system fully compatible with the NCBI taxIDs allowing to create their full or ranked linages.

The mkCOInr tool is a series of Perl scripts necessary to download sequences from BOLD and NCBI, build the COInr database and customize it according to the users’ needs. It is possible to select or eliminate sequences for a list of taxa, select a specific gene region, select for minimum taxonomic resolution, add new custom sequences, and format the database for BLAST, QIIME, RDP classifier.

The COInr database can be downloaded from https://doi.org/10.5281/zenodo.6555985 and mkCOInr and the full documentation is available at https://github.com/meglecz/mkCOInr.

## Introduction

The use of metabarcoding has increased dramatically in the past decade since the technological advances of this method and the continuous reduction of sequencing costs make it accessible for a wide range of studies (Slatko, Gardner, & Ausubel, 2018). Metabarcoding is applied mainly for biodiversity assessment, but it can be used in other fields such as studying interaction networks or understanding animal diets (Compson, McClenaghan, Singer, Fahner, & Hajibabaei, 2020). It is a valuable alternative to morphology-based inventories, since it is applicable for large-scale studies and wide taxonomic ranges (Compson et al., 2020) without the need of direct and time consuming intervention of experts of specific taxonomic groups (Cahill et al., 2018; Erdozain et al., 2019). However, metabarcoding suffers from a series of pitfalls such as the difficulty to estimate the absolute abundance of taxa due to PCR biases, the presence of false positives and negatives and variable taxonomic resolution among taxa and genetic markers. This calls for a careful study design, the use of controls, the careful choice of analytical tools and a critical interpretation of the results (Alberdi et al., 2019).

One of the difficulties of metabarcoding lies in the taxonomic assignation of sequences and the completeness of the underlying reference databases. Methods of taxonomic assignment can be alignment-based relying of sequence similarities detected by BLAST (Altschul et al., 1997) or VSEARCH (Rognes, Flouri, Nichols, Quince, & Mahé, 2016) implemented in different software (Bokulich et al., 2018; Huson, Auch, Qi, & Schuster, 2007) or based on machine learning (Murali, Bhargava, & Wright, 2018; Pedregosa et al., 2011; Wang, Garrity, Tiedje, & Cole, 2007). However, for all methods, the quality of the reference database is crucial (Hleap, Littlefair, Steinke, Hebert, & Cristescu, 2021). Many methods are sensitive to gaps in the taxonomic coverage of the reference database (Hleap et al., 2021), thus the creation of a reference database with the best coverage available is highly needed.

Several different markers can be used for metabarcoding, since each of them are subject to different taxonomic biases and provide different taxonomic resolution (Ruppert, Kline, & Rahman, 2019). The most widespread markers are the ribosomal RNA markers (18S, 28S, 16S), the Cytochrome Oxidase C subunit I (COI) gene and internal transcribed spacer sequences (ITS) (Creer et al., 2016; Porter & Hajibabaei, 2020). Ribosomal RNA markers allow the amplification from a wide range of taxa, and are the most widely used markers for microorganisms (Creer et al., 2016). The choice of the ideal marker is more difficult when dealing with Eukaryotes. Plants and fungal studies most often use ITS markers, since the COI often contains indels of variable size and location and is not sufficiently variable in these groups. In addition, the taxonomic resolution of plant and fungal ribosomal RNA marker is relatively low (Dentinger, Didukh, & Moncalvo, 2011; Yao et al., 2010). For animals, the use of both ribosomal RNA and COI sequences are widespread (Creer et al., 2016). COI marker is known to be sufficiently variable, thus being able to differentiate most animal species (Andújar, Arribas, Yu, Vogler, & Emerson, 2018). The COI was the most sequenced gene at the beginning of the barcoding era, since it is the main maker of the Barcode of Life database (P. D. N. Hebert, Ratnasingham, & deWaard, 2003), and more animal taxa have been barcoded with COI than with any other markers (Andújar et al., 2018). This provides a solid basis for taxonomic assignment of metabarcoding sequences using COI as a marker.

Regularly updated, curated and marker specific databases are available for ITS (UNITE (Nilsson et al., 2019), PLANTiTS (Banchi et al., 2020)) and for rRNA markers (Greengenes (DeSantis et al., 2006), SILVA (Pruesse et al., 2007)). Conversely, COI sequences are deposited to two different major databases, which are not COI-specific: (i) the nucleotide database of NCBI (hereafter NCBI-nt database; Sayers et al., 2022)) and their European (ENA) and Japanese equivalents (DDBJ) are generalist databases without focusing on a taxon or a gene; (ii) the Barcoding of Life Data System (BOLD; (Ratnasingham & Hebert, 2007)) contains barcoding sequences of several markers, but most of the sequences are from the barcoding fragment of the COI gene. Although the data overlap between these databases is considerable, each of them has sequences that are not found in the other database. Therefore, creating a merged database with sequences from both sources is highly desirable.

A major challenge of pooling sequences from different sources into a single database is to homogenize their taxonomic lineages. This step is not trivial due to the presence of homonyms (e.g. Plecoptera is both an insect order and a moth genus), synonyms and misspellings. Therefore, the only clean solution to deal with taxon names is the use of unique taxonomic identifiers (taxID) which are connected to a non-ambiguous, hierarchical system and allow the identification of the lineage for each taxon. Both the NCBI-nt and the BOLD databases use taxIDs, but the two systems are independent from each other, thus they cannot be simply merged. Finding the equivalent taxon names and taxIDs between the two databases call for a careful comparison of taxon names and their lineages in order to match them. However, a further complication arises from occasional incoherencies of taxonomic lineages from different databases (e. g. *Vexillata* genus is a nematode belonging to the Ornithostrongylidae family according to BOLD, but to the Trichostrongylidae family according to NCBI taxonomy), which further complicates pooling of taxonomic information to a single coherent system.

Merging of COI sequences from the NCBI-nt and BOLD has been attempted in different programmes. BOLD_NCBI_Merger (Macher, Macher, & Leese, 2017) uses a very simple method based on identical taxon names, without avoiding the pitfalls of homonyms. MetaCOXI (Balech, Sandionigi, Marzano, Pesole, & Santamaria, 2022) obtains NCBI taxIDs and taxonomic lineages based on ENA flat files, when available. However, when this information is not offered (the sequence is present only in BOLD), NCBI taxIDs are determined by simply matching taxon names to NCBI taxonomy, without checking for homonymy. Furthermore, taxon names not present in NCBI taxonomy do not receive a taxID, and therefore a taxID system is incomplete.

A further difficulty of creating custom (local) databases is sequence downloading from the original sources. NCBI provides different means of accessing data: a whole database can be downloaded via ftp sites, and filtered subsequently, or Application Programming Interfaces (API) are provided for targeted downloads (Kans, 2021). On the other hand, BOLD systems do not provide an easy way to download the whole public dataset, and the use of BOLD APIs needs a considerable optimization to be able to access large datasets. Although bold R package (https://docs.ropensci.org/bold/) is available to download data from BOLD, it is subject to failure for large taxa and takes several hours or days, according to requested data size.

The mkCOInr tool was designed to create the COInr database, which includes all COI sequences from NCBI-nt and BOLD sequences, irrespective of the region of the gene covered and the taxonomic group. All sequences have a taxID, and all taxIDs form a coherent system compatible with, but not limited to, the NCBI taxIDs, allowing to unambiguously obtain taxonomic lineages even for taxon names with homonyms. Sequence redundancy within taxa is eliminated to reduce database size, without losing information. This database is freely available and can be easily and quickly downloaded from https://doi.org/10.5281/zenodo.6555985, thus saving the most complicated and time-consuming steps of custom database creation. Users can customize the downloaded database using mkCOInr scripts and format them to be able to use it with their preferred taxonomic assignment tool. It is possible to add local sequences, select or eliminate sequences of a list of taxa, filtering sequences for minimum taxonomic resolution, and choosing a gene region. The COInr database is planned to be updated annually, but all scripts are available with detailed documentation to re-create it at any time or produce a different database by modifying some of the filtering options.

## Material and Methods

mkCOInr is a series of Perl scripts that can be executed in command line, thus being easily integrated into other pipelines. They were written for Linux OS and can run on MacOS or other Unix environments. The Windows Subsystem Linux (https://docs.microsoft.com/en-us/windows/wsl/) allows Windows users to run mkCOInr scripts. Special care was taken to reduce dependencies to easy-to-install, third-party programmes without the use of special packages. BLAST (Altschul et al., 1997), vsearch (Rognes et al., 2016), cutadapt (Martin, 2011), and NSDPY (R. Hebert & Meglécz, 2022) can all be installed either through the Python Package Index (PyPI) or standard program repositories.

Fig 1 represents a complete flowchart of the pipeline. A tutorial and detailed documentation is available at https://github.com/meglecz/mkCOInr.

**FIGURE 1.**
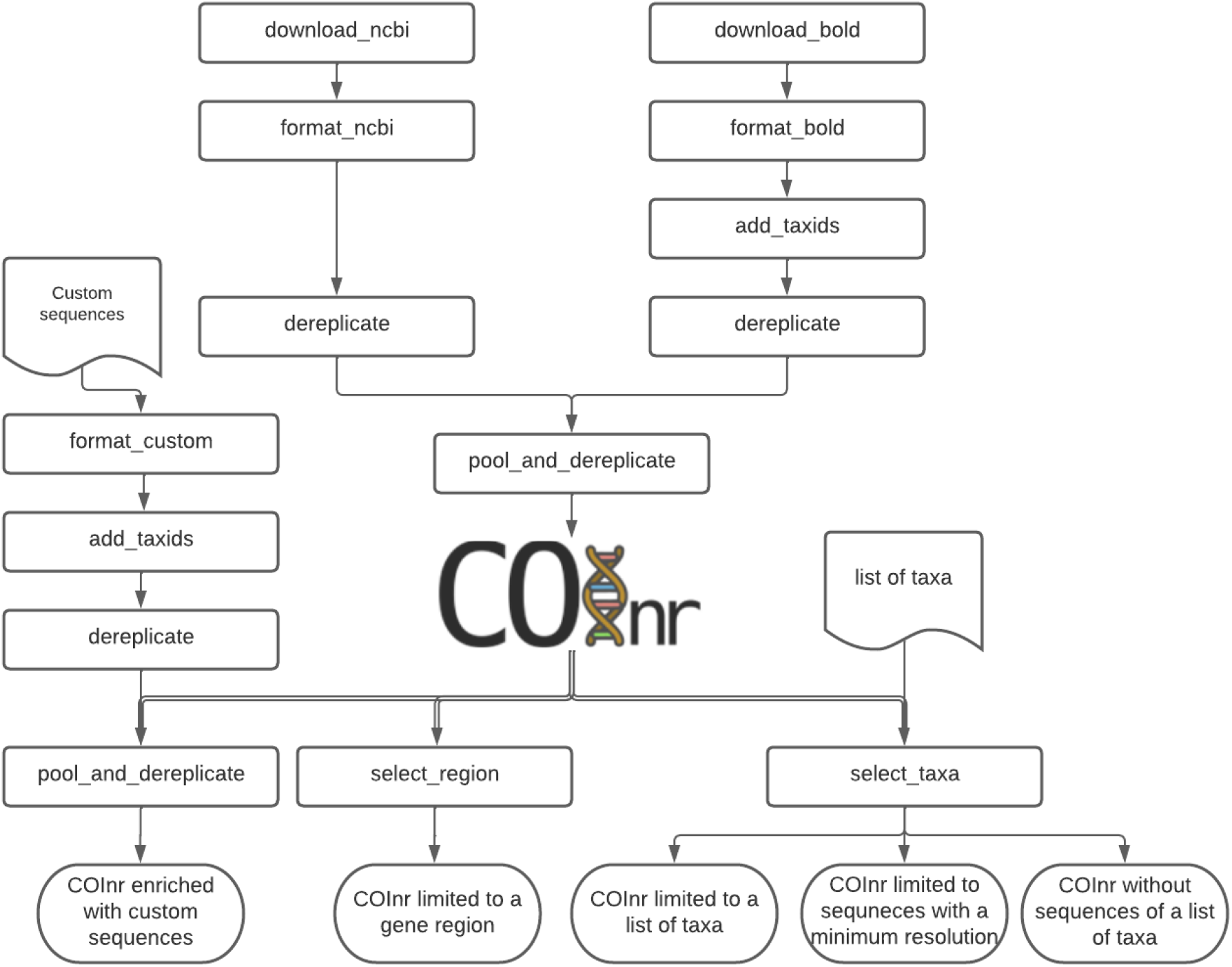
Flowchart of mkCOInr. Double lines represent the different options for customizing the COInr database. These steps can also be consecutive.

### Construction of the COInr database

#### NCBI

NCBI sequences were downloaded with by the NSDPY (R. Hebert & Meglécz, 2022) python package using the following request :

~~~
nsdpy - r “ COI OR COX1 OR CO1 OR COXI OR (complete[Title]
AND genome[Title] AND Mitochondrion[Filter])” - T -v --cds
~~~

This allowed the download of all coding DNA sequences (CDS) returned with the keyword search for COI, CO1, COXI or COX1, and CDS from complete mitochondrial genomes. The scope of this search was intentionally very wide, and the downloaded sequences were further filtered by the *format_ncbi. pl* script to (i) only retain CDS with gene and protein names corresponding to COI, and (ii) eliminate genes with introns and sequences from environmental or metagenomic samples. Sequences with more than five consecutive internal Ns, and outside of the length range of 100-2000 nucleotides were also eliminated. Open nomenclature was not accepted in taxon names. If the taxID did not correspond to a correct Latin name format, the smallest taxon with a correct Latin name in the lineage was chosen for the sequence (e. g. *Acentrella* sp. AMI 1, taxID: 888165, rank: species was replaced by *Acentrella*, taxID: 248176, rank: genus). Sequences were then subjected to taxonomically aware dereplication by the *dereplicate. pl* script. Within each taxID, all sequences that were a substring of another sequence were eliminated. This allows to reduce the size of the database without losing information and keeping intraspecific variability.

#### BOLD

A list of taxa was established from the taxonomy page of BOLD Systems (https://www.boldsystems.org/index.php/TaxBrowser_Home), where each taxon had fewer than 500 000 specimen records. All public sequences of the above list and associated information were downloaded from BOLD, using the *download_bold. pl* script that uses the BOLD APIs. For each taxon, the integrity of the downloaded files and the number of records were checked, and the download was repeated automatically in case of failure. From the raw downloaded files, COI sequences (COI-5P, COI-3P) were selected if they did not contain more than five consecutive internal Ns and were in the length range of 100-2000 nucleotides. As for NCBI sequences, the smallest taxon in the BOLD lineage with a correct Latin name was chosen for the sequence to avoid open nomenclature. All unique lineages were then listed with the corresponding sequence identifiers (sequenceID) and for each lineage a taxID was determined using the *add_taxids. pl* script: the smallest taxon is identified in each BOLD lineage, where the name is matching a taxon name in the NCBI taxonomy database (including synonyms), and at least 60% of the taxon names in the BOLD lineage match the NCBI lineage. For example, for the BOLD lineage of ‘ Chordata, Actinopterygii, Trachiniformes, Pinguipedidae, *Parapercis, Parapercis somaliensis’*, the *Parapercis* genus matches the 215380 NCBI taxID, even if the orders are different in BOLD and NCBI (Trachiniformes and Uranoscopiformes, respectively). In the next step, a taxon under the smallest taxon with NCBI taxID was attributed to an arbitrary, negative taxID, and the new taxID was integrated to the taxID system, with the NCBI taxID as a parent. The newly created taxID was then added to the taxID system and it was characterized by a taxon name, a taxonomic rank and the taxID of its direct parent, forming a hierarchical system. This hierarchical taxID system allows the creation of the lineage of any taxID unambiguously, even in case of homonymy and synonymy. As for NCBI sequences, the filtered BOLD dataset was dereplicated by the *dereplicate*.*pl* script.

To compare the effect of using only correct Latin names (as in COInr) or accepting all taxon names presents in the input databases, the above pipeline was run a second time using systematically the smallest taxon in each lineage, even if it did not correspond to a correct Latin name.

### The COInr database

The BOLD and NCBI datasets were pooled into one single dataset by the *pool_and_dereplicate. pl* script, where sequences for the taxIDs shared by the two source databases were dereplicated, while sequences from taxIDs unique to one of the sources were simply added to the combined database. This database is a starting point to create more specific custom databases according to the users ‘ needs.

The core database consists of two simple-to-parse tsv files (tab separated values). The sequence file has three columns (sequenceIDs, taxIDs and sequences), and contains sequences of all taxonomic groups that can cover any COI region, with variable taxonomic resolution from species to phylum level. The taxonomy file contains taxIDs, scientific names, parent taxIDs, taxonomic rank and taxonomic level index. The taxonomic level index contains integers from 0 to 8 each corresponding to a major taxonomic level (rank): root, superkingdom, kingdom, phylum, class, order, family, genus, species. Intermediate taxonomic levels have 0.5 added to the next major taxon level index (e.g. 7.5 for subgenus). This file allows the reconstruction of the complete lineages of all taxa or the ranked lineages containing only the major taxonomic ranks.

### Customizing the COInr database

The COInr database can be modified according to users’ needs. Sequences can be selected for a list of taxa or on the contrary, removed from the database through the *select_taxa. pl* script. The script will also produce a lineage and a taxID for each taxon in the taxon list, allowing users to check for potential errors due to homonyms. In case of incoherence, the taxon list enriched by the correct taxIDs can be used to rerun the script with more precise selection. The same script also allows selecting sequences with a minimum taxonomic resolution.

The *select_region*.*pl* script trims the sequences to a specific region of the COI gene. Using the usearch_global command of vsearch (Rognes et al., 2016), sequences of the database are aligned to a small, taxonomically diverse pool of the sequences, which have already been trimmed to target region (target_region_fas). The sequences of the core database are trimmed according to the alignment positions. The target_region_fas file can be provided by the users or can be produced by the same script by making an E-PCR on the core database using cutadapt (Martin, 2011).

The COInr database can also be completed by custom sequences. Users will need a taxon name and sequenceID for each custom sequence. The *format_custom*.*pl* script will produce a lineage file for each input taxa, which should be checked, and eventually corrected and completed by the users. The *add_taxids*.*pl* script will add taxIDs to each lineage and complete the input taxonomy file (part of the COInr database). Sequences should then be dereplicated by the *dereplicate*.*pl* script and added to the COInr database using the *pool_and_derelplicate*.*pl*.

Fig 1 represents the customizing options on mkCOInr, each of them starting from the COInr database. However, the different steps can also be successive to produce a final database. For example, it is possible to start by selecting sequences for a list of taxa, then adding custom sequences to the newly created database, which in turn can be trimmed to the target region.

### Format Database

The very simple format of the database (sequence file and taxonomy file both in tsv format) allows users to easily obtain a database in their desired format. The *format_db. pl* script can produce databases ready to use for BLAST, RDP_classifier, and QIIME. The ‘ full’ option will produce a single tsv file with sequence IDs, ranked lineages, taxIDs, and the sequences allowing user to parse, and produce basic statistics on the database content (e.g. number of sequences of each taxon).

## Results

Table 1 summarizes number of taxa and sequences in the initial databases before and after taxonomically aware dereplication, and after pooling and dereplicating sequences from BOLD and NCBI-nt to the COInr database. After the initial quality control, NCBI and BOLD databases contained 3.9 M and 7.6 M COI sequences respectively, belonging to approximately 200 000 taxa with correct Latin names in both databases. Taxonomically aware dereplication within each of the source databases resulted in 1.7 M and 2.8 M nonredundant sequences, corresponding to 58% and 63% reduction in NCBI and BOLD databases, respectively. The total number of taxa was 268 438 after pooling NCBI and BOLD, 69% of which was shared between the input databases, 14% and 17% of unique to NCBI and BOLD, respectively. After pooling the databases and dereplication, 90% of the sequences were from taxa present in both databases, while 4% and 6% specific to NCBI and BOLD, respectively. Overall, the 11.5 M input sequences were reduced to 3.3 M by eliminating redundancy between the two input databases, and within each taxon.

**TABLE 1.** The number of taxa and COI sequences of the input databases (NCBI-nt, BOLD), and in the COInr database (May 2022). COInr is the results of pooling and taxonomically aware dereplication of sequences in the input databases.

|  | N° taxIDs | N° sequences |
| --- | --- | --- |
| After initial quality control |  |  |
| NCBI | 221 565 | 3 920 624 |
| BOLD | 231 425 | 7 590 488 |
| After dereplication within input DB |  |  |
| NCBI | 221 565 | 1 657 602 |
| BOLD | 231 425 | 2 843 248 |
| After pool and dereplicate (COInr) |  |  |
| Shared by BOLD and NCBI | 184 552 | 2 944 524 |
| Unique to NCBI | 37 013 | 124 811 |
| Unique to BOLD | 46 873 | 190 319 |
| **Total** | **268 438** | **3 259 654** |

Apart from sequences of animals, which made 99% of the database and corresponded 97% of the species, other Eukaryotes (plants, Fungi) and even some Bacteria and Archaea sequences were also present in the database (Table 2). Within Metazoa, 83% of the sequences were from Arthropoda that corresponds to 74% of the animal species of the database.

**TABLE 2.** The number of taxa and sequences by phylum.

|  | class | order | family | genus | species | seqN |
| --- | --- | --- | --- | --- | --- | --- |
| Eukaryota |  |  |  |  |  |  |
| **Metazoa** | **126** | **679** | **5 793** | **60 175** | **251 755** | **3 227 851** |
| Arthropoda | 20 | 135 | 2 486 | 41 975 | 185 721 | 2 692 056 |
| Chordata | 14 | 178 | 1 202 | 8 646 | 35 960 | 272 027 |
| Mollusca | 9 | 69 | 649 | 4 213 | 14 860 | 134 996 |
| Annelida | 3 | 27 | 152 | 1 035 | 3 603 | 39 322 |
| Platyhelminthes | 7 | 45 | 231 | 915 | 2 275 | 21 776 |
| Echinodermata | 6 | 47 | 185 | 709 | 1 854 | 19 590 |
| Nematoda | 3 | 20 | 169 | 608 | 1 873 | 14 117 |
| Cnidaria | 7 | 29 | 268 | 896 | 2 474 | 11 212 |
| Rotifera | 3 | 9 | 29 | 78 | 270 | 6 452 |
| Porifera | 5 | 33 | 130 | 412 | 1 147 | 3 707 |
| Nemertea | 4 | 10 | 40 | 120 | 347 | 3 032 |
| Acanthocephala | 5 | 10 | 21 | 62 | 149 | 1 811 |
| Tardigrada | 3 | 7 | 24 | 68 | 234 | 1 615 |
| Bryozoa | 4 | 7 | 69 | 132 | 286 | 1 296 |
| Chaetognatha | 2 | 5 | 10 | 23 | 47 | 1 051 |
| Onychophora | 2 | 2 | 3 | 38 | 111 | 989 |
| Sipuncula | 1 | 5 | 9 | 24 | 74 | 526 |
| Other | 28 | 41 | 116 | 221 | 470 | 2 276 |
| **Viridiplantae** | **30** | **115** | **280** | **990** | **1 834** | **2 362** |
| Streptophyta | 17 | 90 | 235 | 920 | 1 722 | 2 174 |
| Other | 13 | 25 | 45 | 70 | 112 | 188 |
| **Fungi** | **32** | **71** | **147** | **265** | **739** | **1 984** |
| Ascomycota | 13 | 38 | 71 | 139 | 433 | 1 108 |
| Basidiomycota | 8 | 20 | 61 | 105 | 261 | 585 |
| Other | 11 | 13 | 15 | 21 | 45 | 291 |
| **undef** | **55** | **202** | **444** | **1 306** | **4 928** | **26 604** |
| Rhodophyta | 4 | 37 | 130 | 628 | 2 228 | 13 191 |
| Oomycota | 1 | 11 | 18 | 57 | 804 | 3 738 |
| undef | 19 | 69 | 141 | 344 | 834 | 3 685 |
| Apicomplexa | 3 | 5 | 13 | 32 | 351 | 2 951 |
| Ciliophora | 6 | 21 | 60 | 103 | 291 | 1 489 |
| Bacillariophyta | 5 | 24 | 36 | 61 | 206 | 920 |
| Other | 17 | 35 | 46 | 81 | 214 | 630 |
| **Archaea** | **1** | **2** | **2** | **2** | **2** | **2** |
| **Bacteria** | **7** | **14** | **16** | **33** | **46** | **850** |
| **Viruses** | **1** | **1** | **1** | **1** | **1** | **1** |

To evaluate the effect of using non-standard taxon names, corresponding to open nomenclature (e. g. *Allograpta aff. argentipila, Alona guttata group, Macrobiotus cf. hufelandi*) or correct Latin names completed by arbitrary identifiers (e.g. *Macrobathra sp. ACL2485, Abablemma BioLep730, Abacarus sp. GD111*), two databases were created: COInr, where only correct Latin names were used and the all-names database created by the same pipeline, with the exception that all taxon names were accepted regardless of their format (e.g. *Lepidoptera sp. 096 PS-2011* was used as it is instead of the taxID of *Lepidoptera* order). The total number of taxa in NCBI was more than three times higher when using all names (769 956 vs. 221 565). This difference was smaller, yet considerable for the number of BOLD taxa (322 927 vs. 231 425) for the all-names and Latin names databases (Table 3).

**TABLE 3.** Comparison of the number of sequences and taxIDs when accepting all taxon names or using only formal Latin names.

|  | **NCBI** | **NCBI** | **BOLD** | **BOLD** |
| --- | --- | --- | --- | --- |
|  | Latin names | All names | Latin names | All names |
| Total number of sequences | 1 630 665 | 1 768 768 | 2 815 860 | 2 826 583 |
| % of sequences present in different taxIDs | 1,44% | 3,99% | 0,87% | 1,08% |
| Total number of taxIDs | 221 565 | 769 956 | 231 425 | 322 927 |
| % of taxIDs sharing sequences with another taxIDs | 9,80% | 28,91% | 10,97% | 13,21% |
| % of taxIDs without unique sequences | 1,82% | 25,45% | 1,57% | 5,59% |

The proportion of the identical sequences shared by different taxa was also higher, when accepting all taxon names compared to using only Latin names, especially for NCBI: 4.0% vs. 1.4% for NCBI, 1.1% vs. 0.9% for BOLD. Similarly, the proportion of taxIDs sharing identical sequences was higher using all names: 28.8% vs. 9.8% for NCBI, 13.2% vs. 11.0% for BOLD. The same tendency was observed for the proportion of the taxIDs that had only sequences identical to other taxa: 25.5% vs. 1.8% for NCBI, 5.6% vs. 1.6% for BOLD (Table 3).

## Discussion

The need for high-quality database can be measured by the number of published databases and methods of their construction. Several tools exist such as the CRUX database Builder integrated to Anacapa (Curd et al., 2019), Metataxa2 Database Builder (Bengtsson-Palme et al., 2018), MetaCurator (Richardson, Sponsler, McMinn-Sauder, & Johnson, 2020), BCdatabaser (Keller et al., 2020), which are not marker-specific. The MIDORI database (Leray, Ho, Lin, & Machida, 2018; Machida, Leray, Ho, & Knowlton, 2017) contains mitochondrial sequences of 13 protein-coding genes. All the above-mentioned databases and tools are based exclusively on NCBI databases or on a dataset already containing a coherent system of lineages. Several COI-specific databases have also been published and are often limited to a target taxon or geographical region. The Eukaryote CO1 Reference Set For The RDP Classifier (Porter & Hajibabaei, 2018) is specifically designed for the RDP classifier and focuses on Arthropoda and Chordata. It contains NCBI and BOLD sequences of at least 500 bp, but the last update is from 2019 and the scripts for re-creating the database are not available. The Meta-Fish-Lib (Collins et al., 2021) is a generalized, dynamic reference library fishes. MitoFish (Sato, Miya, Fukunaga, Sado, & Iwasaki, 2018) is limited to fish mitochondrial sequences. The MARES database (Arranz, Pearman, Aguirre, & Liggins, 2020) is specific to marine sequences from BOLD and NCBI. The pipeline is provided to create a new database specific to the users’ needs. However, a potential source of problems for installing and using the scripts is the high need of third-party programs and packages. METACOXI database (Balech et al., 2022) is a COI database that satisfies many criteria. It includes all Metazoan COI sequences from BOLD and NCBI (ENA) and uses NCBI taxIDs wherever possible. However, for BOLD-specific sequences without NCBI/ ENA accession number, taxIDs are established by simply matching the taxon names without checking for homonymy. Furthermore, taxon names not present in NCBI taxonomy do not receive a unique taxIDs, therefore the database lacks a coherent taxIDs system allowing to avoid all taxonomic ambiguities.

### Use of accepted Latin names

Both BOLD and NCBI contain a high number of taxon names at a species level, with unique taxIDs, which do not correspond to the binomial nomenclature. In most cases they correspond to taxon names of a higher level completed by an identifier or simply completing the taxon name by ‘ sp.’. In principle, they could be proxies of species, but according to my findings, it is unlikely for most cases. When accepting all names as they appear in the input database, a high proportion of the COI sequences are shared between taxa, and most importantly a high proportion of taxa contain only sequences that are identical to sequences of other taxa. COI is known to be variable among most species (P. D. N. Hebert, Cywinska, Ball, & deWaard, 2003) and often shows considerable intraspecific variability (Ratnasingham & Hebert, 2013). The high proportion of shared sequences between taxa suggests that many of the taxa do not correspond to distinct species, but they are the results of an unjustified over-splitting. This phenomenon is particularly pronounced in NCBI, where many abusive examples are found. For example, many genus names in NCBI are completed by the sampleID of BOLD and used as species names (e. g. Platynothrus sp. BIOUG14078-H10): many of them share identical sequences, and do not even correspond to BOLD BINs (Barcode Index Numbers) which would provide some ground for species delimitation. Since the METACOXI database accepts all taxon names as they appear in BOLD or NCBI, it artificially inflates the number of taxa, which are in most cases uninformative to users, hindering efficient, taxonomically aware reduction of redundancy. The COInr database uses only taxa with correct Latin name format. To avoid the loss of sequences, sequences with incorrect taxon names are attributed to the lowest taxon in the lineage with a Latin name. Therefore, sequences are kept in the database, with a conservative level of taxonomic information resulting in a more efficient dereplication, and thus a smaller database without the loss of crucial information.

### Selecting the target region

The COInr database includes sequences that can cover any region of the COI gene. For taxonomic assignment methods based on sequence similarity (Clemente, Jansson, & Valiente, 2011; Huson et al., 2007; Kahlke & Ralph, 2019; Wood & Salzberg, 2014) the database can be used as it is, since sequences of the non-target region will not be returned by BLAST or other similarity searches. The only disadvantage would be the database size, which could be eventually reduced by selecting only the region of the sequence that cover the target region. On the other hand, for taxonomic assignment based on sequence composition or phylogeny (Murali et al., 2018; Nguyen, Mirarab, Liu, Pop, & Warnow, 2014; Rosen, Reichenberger, & Rosenfeld, 2011; Wang et al., 20 07), it is preferable to trim sequences to the target region. This can be done using the mkCOInr tool. It is possible to select only full-length sequences covering the whole target region. However, this comes at the price of losing partial sequences, and thus some taxa. Therefore, mkCOInr can also select sequences that cover user-defined portion of the target region to increase taxonomic coverage.

### Selecting the target groups

Using a large database with a wide taxonomic scope is convenient for users analysing different datasets with a varied taxonomic origin, since the same database can be used and can give a good first approximation of taxonomic assignment of sequences. It can also be helpful to detect contaminant sequences that are not expected in the study (e. g. human sequences or model species studied in the same lab) or sequences outside of the target group of the study (e.g. bacteria, algae, fungi when focusing on animals). By using a generalist database, these sequences can be identified and eliminated. On the other hand, the presence of reference sequences from taxa not relevant to the study can also have disadvantages: the database size is higher and therefore the speed of taxonomic assignment is lower with generalist databases. Moreover, sequences can be assigned to unexpected taxa if the taxonomic coverage of the target group is incomplete. This can be avoided with databases specific to the target group (Axtner et al., 2019; Mathon et al., 2021; Valentini et al., 2016). For example, many sequences from marine samples can be erroneously assigned to insects when using a generalized database, which is the combined result of the facts that most marine groups are insufficiently covered in the reference databases (Mugnai et al., 2021), and an overwhelming majority of the sequences are from insects (73%). Therefore, the possibility to easily create custom databases specifically tailored to the users’ needs is particularly important, and the mkCOInr provides the necessary tools to make this selection.

### Selecting sequences with different taxonomic resolution

Another consideration when creating custom databases is whether to keep reference sequences with incomplete lineages. Most sequences of a reference database assigned to an insect order without further precision is likely to be useless, since most insect reference sequences are determined at least to the genus level, and the taxonomic coverage of this group is wide. On the contrary, for less well-covered groups, especially if species or higher-level groups are difficult to identify morphologically (e.g. Nematoda, Rotifera), reference sequences with partial lineages are still informative.

### Database curation

Erroneously annotated sequences in the reference database can have serious consequences on taxonomic assignations. Ideally, a reference database should be curated to identify incorrectly assigned sequences. Unfortunately, both NCBI and BOLD databases contain mislabeled sequences. Published methods aiming to curate databases are not applicable to large databases, since either the run time would be prohibitive or include a manual step for the curation (Collins et al., 2021; Kozlov, Zhang, Yilmaz, Glöckner, & Stamatakis, 2016; Rulik et al., 2017). The COInr database is too large to be able to run a curation step, which should be kept in mind when using the full database. However, if a small custom database is created from COInr, this curation step becomes feasible and strongly recommended.

## Conclusions

The COInr database can be used for taxonomic assignations of COI sequences as it is, since it is not limited in its taxonomic scope, or to a particular region on the gene. It is also a good starting point to create local, custom databases, since it saves the most time-intensive and complicated steps of database creation: (i) downloading a large number of sequences (ii) creation of a coherent taxID system to avoid ambiguity due to homonymy and synonymy (iii) and sequence dereplication.

The mkCOInr package provides the necessary tools to both to re-create a whole COInr database, between the planned annual updates, and produce custom database starting from COInr. The possibility of refining the taxonomic composition of the database, selection of the gene region and formatting the output to widely used database formats (blast, rdp, qiime) are filling the need for an easy way of creating customized COI databases.

## Acknowledgements

I thank Francesco Mugnai for testing mkCOnr and making valuable comments on their use, documentation and the paper and Gabriel Nève for language editing.

## Data Accessibility and Benefit-Sharing

The complete COI database can be downloaded from https://doi.org/10.5281/zenodo.6555985. All scripts are available in https://github.com/meglecz/mkCOInr including full documentation.

## Author Contributions

EM has designed the research, wrote the scripts, analysed the data and wrote the manuscript.

